# 3D clustering analysis of super-resolution microscopy data by 3D Voronoi tessellations

**DOI:** 10.1101/146456

**Authors:** Leonid Andronov, Jonathan Michalon, Khalid Ouararhni, Igor Orlov, Ali Hamiche, Jean-Luc Vonesch, Bruno P. Klaholz

## Abstract

Single-molecule localization microscopy (SMLM) can play an important role in integrated structural biology approaches for example at the interface of cryo electron microscopy (cryo-EM), X-ray crystallography, NMR and fluorescence imaging to identify, localize and determine the 3D structure of cellular structures. While many tools exist for the 3D analysis and visualisation of crystal or cryo-EM structures little exists for 3D SMLM data which can provide fascinating insights but are particularly challenging to analyze in three dimensions especially in a dense cellular context. We developed 3DClusterViSu, a method based on 3D Voronoi tessellations that allows local density estimation, segmentation & quantification of 3D SMLM data and visualization of protein clusters within a 3D tool. We show its robust performance on microtubules and histone proteins H2B and CENP-A with distinct spatial distributions. 3DClusterViSu will favor multi-scale and multi-resolution synergies to allow integrating molecular and cellular levels in the analysis of macromolecular complexes.

---

Structure-function studies of macromolecular complexes are increasingly moving towards cellular structural biology which requires developing multi-resolution and multi-scale correlative approaches to address the molecular and cellular organization of living cells^1,2^. The strength of such strategies is illustrated for example by correlative light and electron microscopy (CLEM) approaches^3–7^. Methods such as protein X-ray crystallography, NMR and cryo electron microscopy (cryo-EM) have had a strong impact over the past decades thanks to their ability of analysing and visualising macromolecules directly in 3D. These well-established 3D techniques encompass dedicated software tools to analyze and also visualize 3D structures, while no equivalent tools exist up to now in the field of single-molecule localization microscopy (SMLM). SMLM includes techniques such as stochastic optical reconstruction microscopy (STORM^8^) or photo-activated localization microscopy (PALM^9^) and gives access to the properties of every single fluorescent molecule, which allows precise determination of their lateral (X,Y) coordinates^10,11^, separation of spectrally close fluorophores^12^ and single-molecule FRET^13^, but also the possibility of determining the axial (Z) position of the fluorophores with sub-diffraction precision. Methods for 3D SMLM include bi-plane detection^14^ or modifications of the point spread function (PSF) through either astigmatism^8^ or double-helix PSF^15^, or the 4Pi optical setup^16^. Recently, it has also became possible to get some sub-diffraction 3D information from 2D data using a photometry approach^17^. Despite these experimental possibilities for the acquisition of 3D SMLM data, methods for processing these in 3D are not as well developed. For 2D SMLM data, means for visualization^18,19^, co-localization analysis^19,20^ and segmentation^21–24^ have been reported including processing methods based on tessellations, namely on 2D Voronoi diagrams and Delaunay triangulations, which have advantages over other approaches because they allow for efficient visualization^19^, unambiguous local density estimation^19,25^, noise reduction and multi-scale segmentation^25^. Even though most of these approaches are described as potentially extendable to the 3D case, to date only DBSCAN^26,27^ and Getis and Franklin's local point pattern analysis have been used for 3D segmentation of localization data^28,29^; other methods such as Bayesian clustering^23^ and Voronoi-based segmentations in fact remain 2D approaches as neither SR-Tesseler nor ClusterVisu can process 3D data^12,18^.

Processing data directly in 3D rather than in 2D turns out to be crucial to avoid artefacts due to the enhanced complexity in 3D as compared to the 2D case, especially in a dense cellular context. An analysis in 2D would lead to an overlap between neighboring structures which are otherwise separate in 3D space. Using test data with clusters distributed randomly in a 3D space with noise, we show that a simple 2D clustering analysis of the data can lead to unprecise results with erroneous size estimations (Fig. 1). With increasing density of clusters and of background noise, the point distributions obtained for a 2D analysis become close to random due to the overlay of 3D structures in a 2D image (Fig. 1B–C), while 3D segmentation of the same data is still robust and retrieves the clusters properly (Fig. 2C-H). This suggests that in order to avoid information loss and instead precisely determine properties of 3D objects which are labelled within a cell or a tissue, it is essential to acquire and also segment and represent the data in three dimensions, especially in crowded environments such as chromatin. In the following we describe the concept of three-dimensional Voronoi-based cluster analysis which has never been applied to 3D SMLM data before. We evaluate the performance of this method with synthetic and experimental data and provide examples of visualization and quantitative analysis of 3D SMLM data.

**Figure 1.**
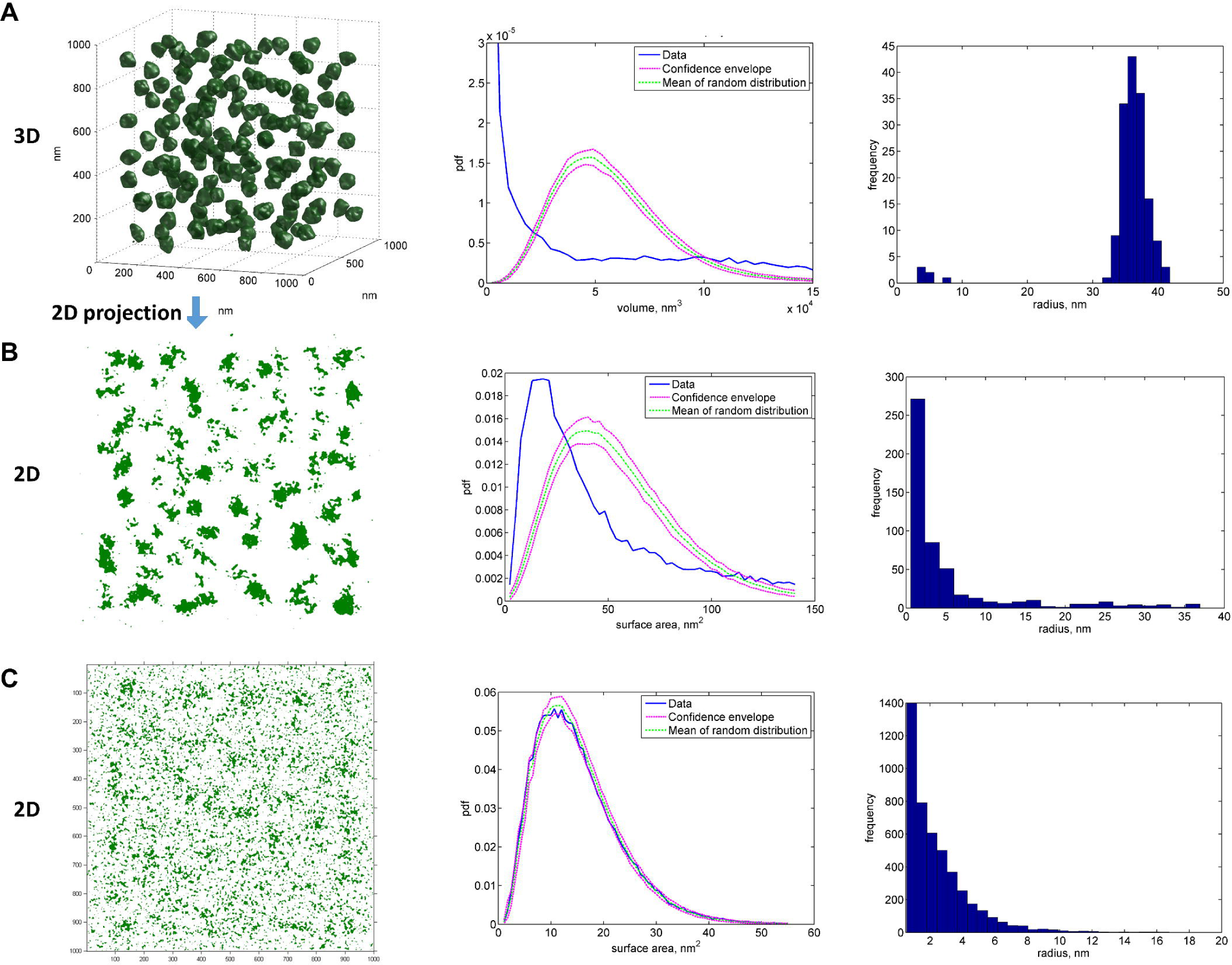
Importance of 3D cluster analysis of localization data. (**A**)Clusters in a 3D dataset (number of clusters = 150, density of localization within clusters = 6-10^−4^ nm^−3^, in the background = 8-10^−6^ nm^−3^) can be easily segmented using the 3D Voronoi diagram approach described in the current work. (**B**) The same dataset projected on the X-Y plane and processed with a 2D segmentation method^19^ does not allow to determine the correct shape, size and number of clusters. (**C**) Similar dataset with higher background density (6-10^−5^ nm^−3^), lower cluster density (4-10^−4^ nm^−3^) and higher number of clusters (175) becomes undistinguisheble from a random distribution when projected in 2D, but still can be successfully processed in 3D (see later Fig. 2C–H). In the left column are the clusters determined with the 3D Voronoi approach described in the current work (A), with our previously reported 2D Voronoi approach^19^ (B) and using the mean local density as the threshold for binarization of 2D data^25^ (C). In the middle column are probability density functions of volumes (A) or surfaces (B, C) of Voronoi cells for the processed datasets (blue) in comparison with that of the background model, a spatially randrom distribution (magenta and green). The experimental curve inside the confidence envelope (C) indicates that the experimental dataset is undistinguishable from the background model and so no clusters can be detected with confidence. In the right column are histograms of the equivalent radius of the detected clusters, to be compared with the simulated cluster radius of 30 nm.

A Voronoi diagram for the 3D case is a space partitioning into polyhedron regions such that every polyhedron would be the locus of points closest to the corresponding seed^30^. Similarly to the 2D case where each event forms a seed (Fig. 2A), in 3D the seeds are defined by the single-molecule (X,Y,Z) coordinates, while the Voronoi polyhedrons can be considered as the regions of influence of the corresponding fluorophore molecules (Fig. 2B). Because the geometrical properties of Voronoi regions reflect the environment characteristics of a given molecule, we can use their short-range collective properties determined from the diagram in addition to the experimentally determined coordinates and intensities of the fluorophores. One of the most important properties, the density *d* in the neighborhood of the molecule i, can be determined simply as the inverse value of the volume *V* of the Voronoi cell: *d_i_ = 1/V_i_*. To segment a SMLM dataset, every localization should be assigned to a cluster or to the background. The minimal local density of molecules inside clusters, *i.e.* the segmentation threshold, can be determined automatically by comparing the distribution of Voronoi cell sizes of the experimental distribution with that of the background noise model, as has been recently shown by us for the 2D case^19^ (practically, the threshold is determined in a region of interest such as the nucleus and then applied to the entire image). Based on Monte-Carlo simulations using an a priori background model (see methods), this type of analysis also indicates whether the experimental distribution is significantly different from the noise model, *i.e.* if it is clustered or not.

**Figure 2.**
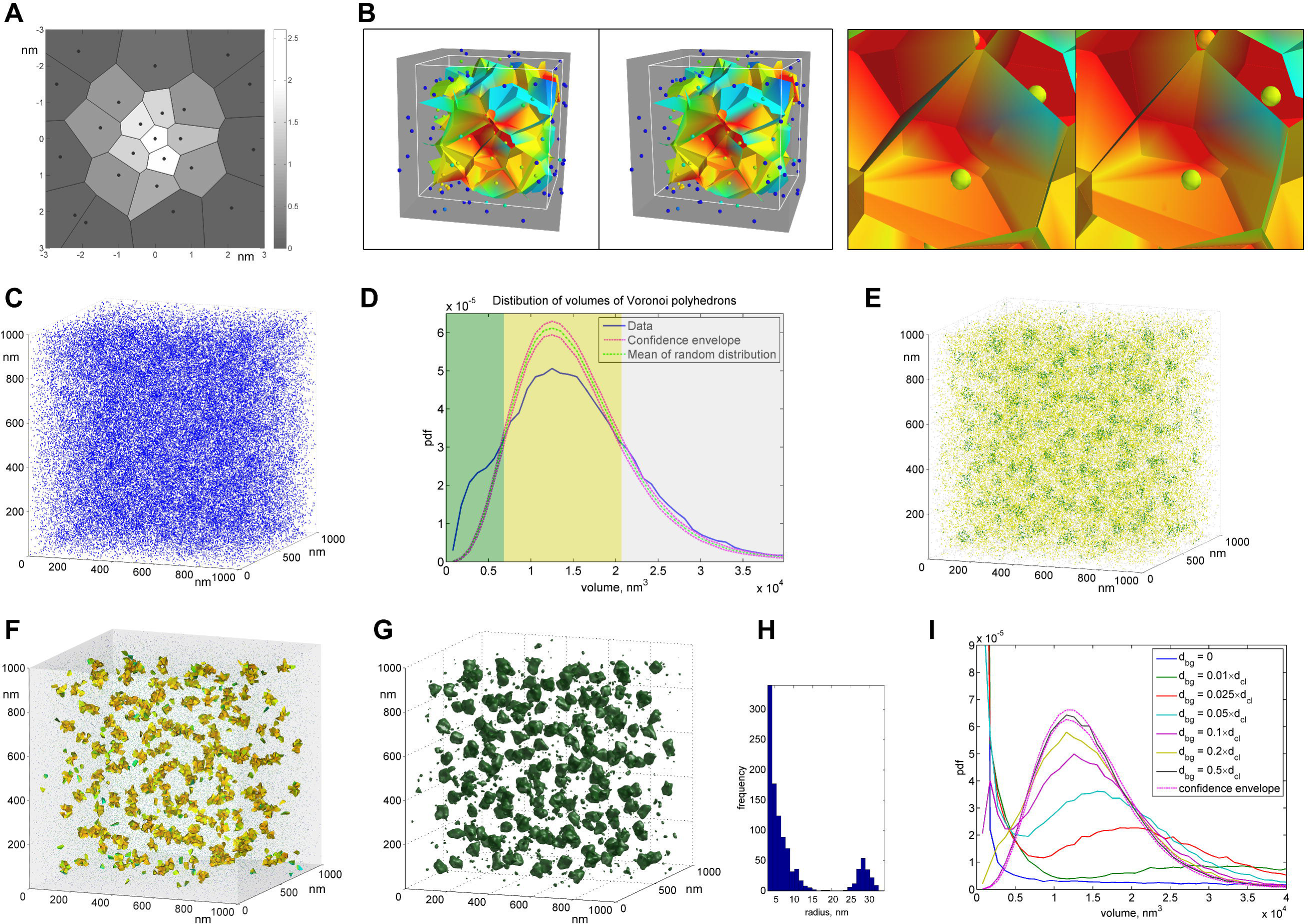
Concept of 3D Voronoi-based segmentation of SMLM data. (**A**)Concept of two-dimensional Voronoi diagrams; the brightness of the polygons is proportional to the local density of points, determined as the inverse value of their surface area. (**B**) The concept of three-dimensional Voronoi tesselations highlights the strong increase in complexity upon transition from 2D to 3D data (stereo representations; zoomed region on the right; small spheres indicate the coordinates of the fluorophores; Voronoi diagram cells are colored according to their size, from blue (big cells) to red (small cells) and allow deriving the number of molecules present in a given 3D cluster with associated radius and volume values; 3D visualization tool available under http://cbi-dev.igbmc.fr/cbi/voronoi3D/tree/master/visualisation). (**C**) Simulated 3D volume of clustered points in a dense background. (**D**) Distribution of volumes of Voronoi polyhedrons of the clustered dataset (blue); mean values (green) and confidence envelope (red) of a set of distributions for datasets with randomly placed points obtained from Monte-Carlo simulations. The three characteristic regions: small clustered polyhedrons (green), intermediate and huge polygons corresponding to background (yellow and gray). (**E**) The original points displayed in colors accordingly to the three regions allow to visually delimit the clusters. (**F**) Small Voronoi polyhedrons correspond to the clusters. (**G**) Density map binarized at the level of the determined threshold allows cluster analysis. (**H**) Histogram of the equivalent radius of the clusters. The first peak corresponds to the small clusters originated from fluctuations in the dense background, the second peak with radius of (28.5±1.6) nm reveals the simulated clusters. The dataset on (**C-H**) consists of 175 clusters with a radius of 30 nm. The density of molecules inside the clusters is 4.10^−4^ nm^−3^, the density of background localizations is 6-10^−5^ nm^−3^. (**I**) Distribution behavior of volumes of Voronoi polyhedrons at different noise and cluster density ratios. The dataset contains a fixed total number of points (7.10^4^) and a fixed number of clusters (150) with fixed radius (30 nm).

Using synthetic data to illustrate the 3D Voronoi tessellation concept (Fig. 2C; methods), we show that by randomly distributing the same number of points through the same-sized region we obtain a test dataset with Voronoi polyhedrons that should be compared with those of the clustered dataset. This allows determining three characteristic regions which lie between the intersections of probability density functions from the distributions of Voronoi volumes of the two datasets (Fig. 2D). The smallest polyhedrons correspond to high-density regions that form clusters, while average-sized and big polyhedrons correspond to low-density regions, *i.e.* the background, as highlighted by the color-coded display of the points (Fig. 2E) that can be represented as a Voronoi diagram (Fig. 2F). Next, interpolation of the local densities to a regularly spaced grid generates a 3D density map that can be segmented using the above determined threshold (Fig. 2G) to provide clusters with deterministic shape and properties such as their radius and number of molecules within a given cluster (Fig. 2H, **Fig. S1C**). The example shows many very small clusters that originate from the fluctuations of the local density in the background (Fig. 2H, first peak) and bigger clusters that correspond to the simulated clusters (Fig. 2H, second peak). The equivalent radius of these clusters (28.5±1.6 nm) matches that of their original radius (30 nm) and the number of events inside (36±6) compares well with the simulated number of localizations (45±3). At densities between d_bg_ = 0.1·d_cl_ and d_bg_ = 0.01·d_cl_ (ratio of clustered and noise events) the Voronoi volume distributions break up into two peaks which correspond to the cluster and the background regions, respectively (Fig. 2I). The curve of the completely randomly distributed points crosses the other curves near the valley between the two peaks. This confirms the reliability of our automatic threshold calculation method. For a constant density of clustered molecules, the threshold decreases with increasing background density. This indicates that the cluster volume, which can be determined with confidence, decreases, while it also reduces the number of spurious clusters detected in dense background regions (**Figs. S1 A-C**).

Visualization of 3D data in the form of a 2D image can be difficult, especially in crowded environments with dense fluorophore occurrence due to the missing information in the Z-direction. The example of microtubules (Fig. 3) shows that the scatter plot is useful only for very low density data, otherwise the markers overlap and hide details (Fig. 3A). When displayed as slices of a 3D histogram, the density of molecules is usually not sufficient to ensure good continuity of the images (Fig. 3B). Moreover, when molecules are displayed as Gaussian clouds with a width dependent on the localization precision^29^ the resolution of the image can be impaired^18^. To address these issues, it is useful to display 3D density maps (calculated from the local densities) as slices (Fig. 3C), as projections or as isosurfaces (Fig. 2G; see also later Fig. 4 panels **B & E**). Thanks to the properties of Voronoi polygons and after an interpolation step, the density value at any given voxel takes into account the distribution of molecules in a given 3D region around this voxel and thus ensures a good continuity of the image. Additionally, the Voronoi density maps can be conveniently viewed and analyzed with standard tools for processing confocal microscopy or electron microscopy maps because the local density of fluorophores in localization microscopy is proportional to the brightness in classical microscopy. Another advantage of the method is noise reduction when keeping only the points with a density over the threshold determined from the Monte-Carlo simulations (Figs. 3D-F). The resulting β-tubulin dataset demonstrates a strong reduction of background noise without affecting the densely labelled microtubules’ even when displayed as a 2D histogram image (Figs. 3E&F).

**Figure 3.**
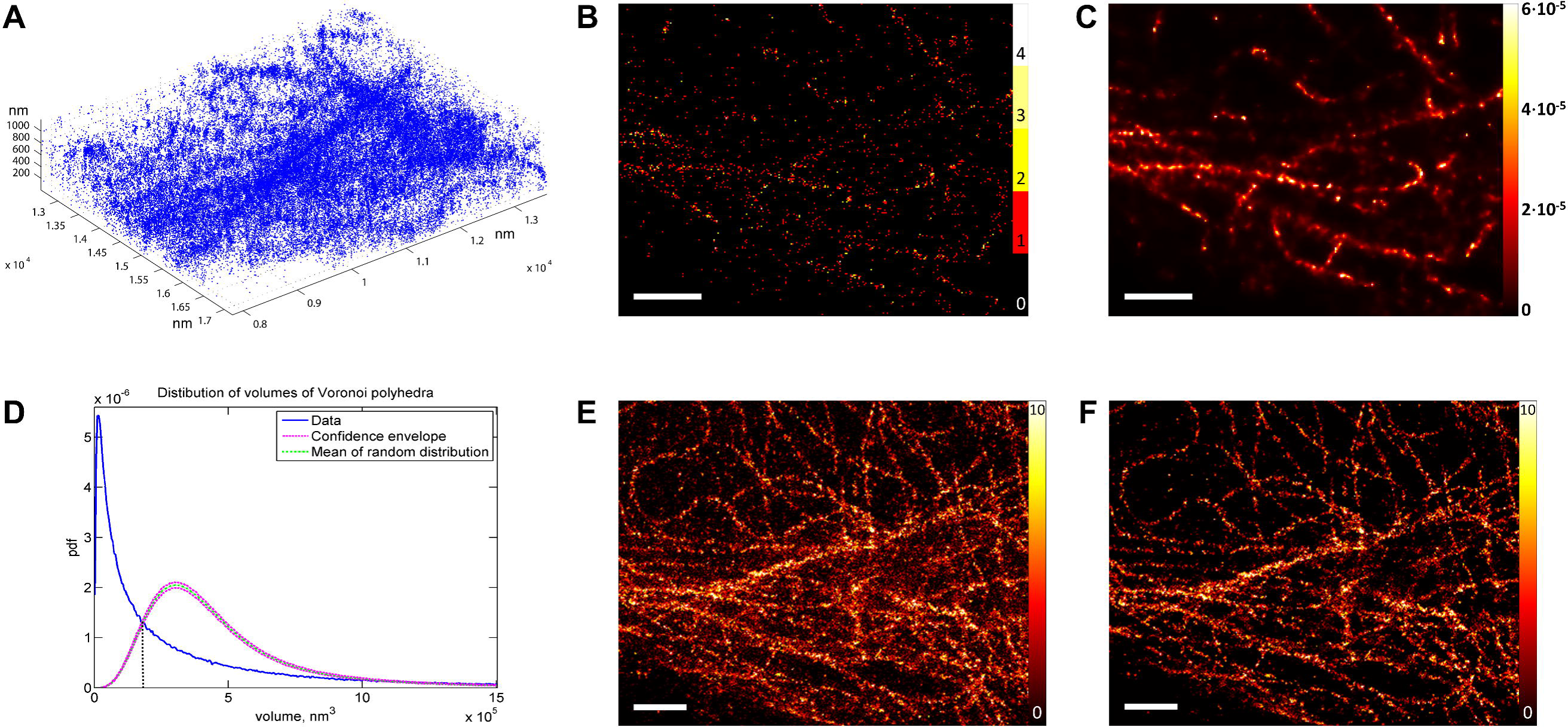
Application of the 3D Voronoi diagram method for visualization and noise reduction on the example of β-tubulin. (**A**) Plot of fluorescent molecules in the form of points (scattered plot) does not allow for correct visualization of dense regions (β-tubulin detected with Alexa-647-labelled secondary antibodies). (**B-C**) A slice of the 3D volume shows more information content when calculated as a Voronoi density map (**C**) compared to a 3D histogram (**B**) (both panels show voxel-thick slices). (**D-F**) Distribution of volumes of Voronoi polyhedrons (**D**) allows noise reduction. (**E-F**) XY projections of the original dataset (**E**) and of the dataset containing only the points with the density over the threshold (**F**). Scale bars: μm. The color bars indicate number of localizations inside pixels for the histogram images (**B, E-F**) or the local density measured in nm^−3^ for the density map (**C**).

We applied the 3D Voronoi segmentation concept to the cluster analysis of two histone proteins, H2B and CENP-A with distinct spatial organizations (Fig. 4). H2B is known to be located inside the nucleus in the form of heterogeneous nanodomains with variable density at euchromatin and heterochromatin regions^31^. The distribution of Voronoi cells of an H2B-labelled dataset lies outside the confidence envelope for the noise model indicating statistically significant clustering (Fig. 4A). Segmented density maps of H2B show bigger and denser clusters at the periphery of the nucleus as expected for heterochromatin (Fig. 4B), the distribution of H2B cluster sizes being rather broad with a median diameter of 42.4 nm (Fig. 4C) in line with previously reported values^31^. In contrast, the histone variant CENP-A that replaces the canonical histone H3^24^ localizes exclusively at centromeric regions^32^ where it plays a key role in determining chromosome location and segregation^23,25^. Our analysis of SMLM CENP-A data shows that the distribution of Voronoi volumes is very different from that of the noise model and that the vast majority of the Voronoi cells is small compared to randomly distributed points indicating a strong clustering (Fig. 4D). Interestingly, we find that CENP-A forms clusters with relatively homogeneous sizes considering the large supra-structure of the centromeric region in the cell (diameter of 260 ± 54 nm, containing an average of 418 localizations within a cluster; **Figs. 4D–F**). The 3D Voronoi segmentation analysis provides for the first time evidence that histone variant CENP-A forms defined clusters in human cells and quantifies their size and occurrence.

**Figure 4.**
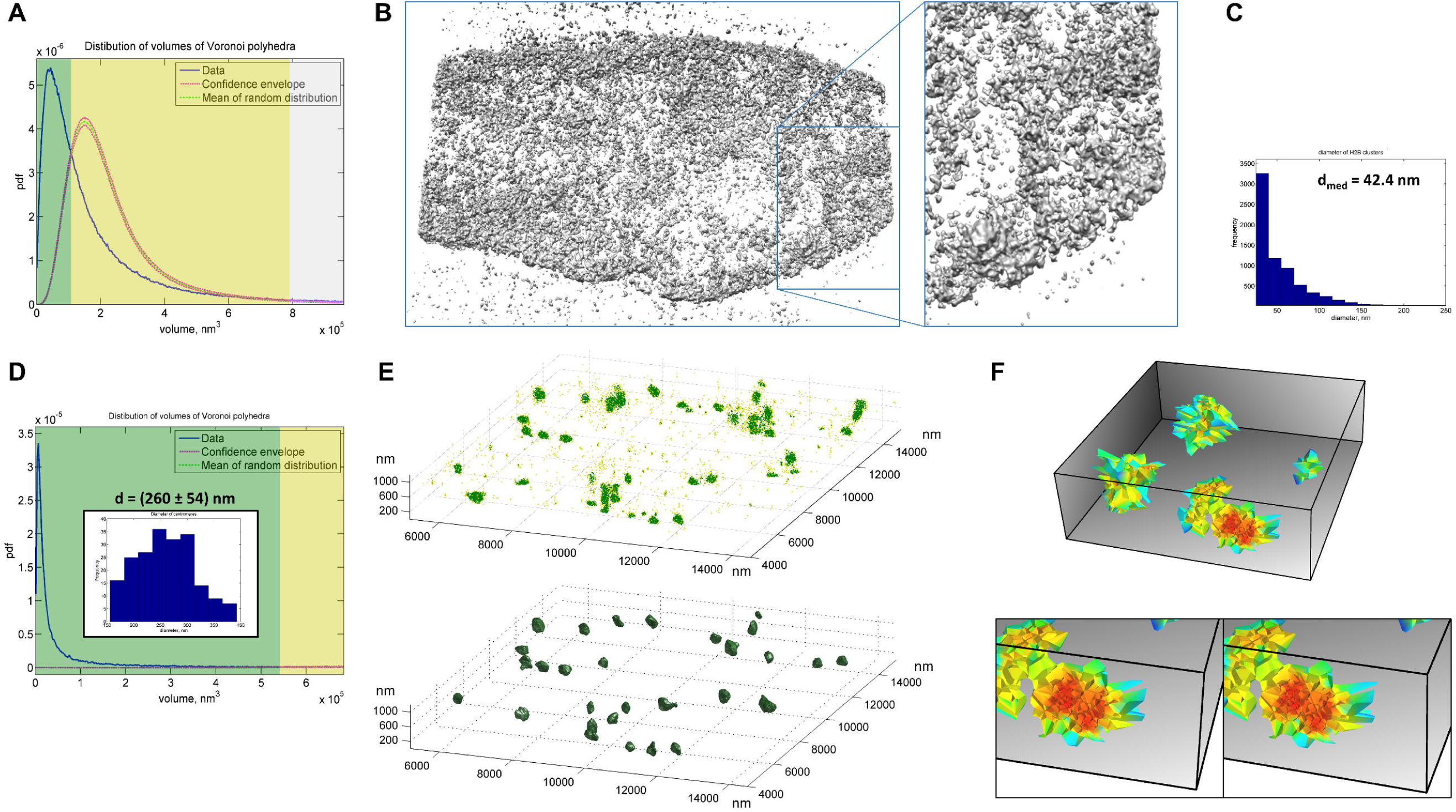
Cluster analysis of histone proteins H2B and CENP-A by 3D Voronoi tesselation. (**A-C**) Histone H2B detected with Alexa-647-labelled secondary antibodies in a HeLa cell. (**A**) The distribution of volumes of Voronoi polyhedrons indicate clustering of histone H2B (color code as in Fig. 1D). (**B**) 3D density map displayed as an isosurface at the level of the threshold in the 3D software Chimera^26^. (**C**) Histogram of the equivalent diameter of H2B clusters with the median value of 42.4 nm. (**D-F**) CENP-A detected with Alexa-647-labelled secondary antibodies in U2OS cells at late G1 phase. (**D**) The distribution of volumes of Voronoi polyhedrons indicate strong clustering of CENP-A which allows quantification of their properties. The histogram of the diameters of the centromeres is shown in the inset. (**E**) Scattered plot of molecules colored according to the local density (top; color code as in panels **A** & **D**); segmented density map displaying CENP-A clusters in the centromeric regions of the cell (bottom). (**F**) 3D representation of CENP-A clusters displayed as Voronoi tessellations in our 3D visualization tool (bottom: zoomed region in stereo representation; mean diameter of 260 nm containing an average of 418 localizations in a given 3D cluster).

Taken together, our novel cluster analysis method introduces a new concept for 3D (rather than 2D) segmentation of SMLM data based on the elegant mathematical properties of 3D Voronoi diagrams that allows deriving 3D cluster radius and volume values and associated molecule numbers. It provides automatic threshold determination and noise filtering. In addition, to avoid distortion effects due to non-isotropic distribution of experimental data points we introduced weighting in Z-direction (see methods; taken into account also for Monte-Carlo simulations). As an example, it takes 6 hours to segment a large experimental dataset with a volume of ~200 μm^3^ that contains 10^6^ localizations using an i7 quad-core single processor with 32 Gb of memory. The calculation includes 50 rounds of Monte-Carlo simulations and interpolation to a 3D density map with a voxel size of 20 nm. Finally, as compared to a 3D implementation of DBSCAN^27^ (see methods) which requires multiple trials through individual runs to optimize parameters manually without additional *a priori* knowledge, our method proceeds in a single run with automatic threshold determination from Monte-Carlo simulations using an a priori background model (into which re-localization events can be included if known) and it proves to be more robust with increasing background densities thanks to noise modelling (**Figs. S1B-D; S1B** includes a statistical analysis of the results, the estimated cluster sizes can vary slightly depending on the background / cluster density). The Voronoi-based method was validated using simulated and experimental data’ illustrating its applicability to different biological objects of interest including cases that are difficult to analyze such as fine chromatin structures in a dense cellular context and larger cellular structures such as tubulin. 3DClusterViSu will be of broad interest to the scientific community as it can be applied to any 3D SMLM data to analyze or re-analyze and quantify 3D information such as spatial organization, local density, volume and shape of labelled protein clusters in 3D and number of molecules within a given cluster. Strikingly, it allows for the first time SMLM analysis and also visualization of macromolecules directly in 3D like in the neighbor fields of cryo-EM and X-ray crystallography; therefore, it will be also a valuable tool for CLEM in integrated cellular structural biology approaches. Standalone software for 3D data processing with a GUI (3DClusterViSu) and a standalone Python-based 3D Voronoi visualization tool with a GUI are available under http://cbi-dev.igbmc.fr/cbi/voronoi3D/repository/archive.zip.

## Author contribution statement

L.A. wrote the software and performed imaging, L.A., I.O, J-L.V. and B.P.K. worked on the concept, J.M. wrote the visualization software, L.A. performed GSDIM experiments, K.O. performed CENP-A labelling, L.A., J-L.V., A.H and B.P.K. analyzed the data, and L.A. & B.P.K. wrote the manuscript. All authors reviewed the manuscript.

## Acknowledgements

We thank Leica Microsystems for the access to the Leica SR GSD 3D system and Didier Hentsch, Pascal Kessler and Yves Lutz of the IGBMC Imaging Centre for support. This research was made possible by the adaptive optics plug-and-play accessory MicAO 3D-SR for PALM/STORM microscopes of Imagine Optic (www.imagine-optic.com). This work was supported by the INCa programme and the Centre Nationale pour la Recherche Scientifique (CNRS). The super-resolution microscope setup is supported by the Alsace Region and by the French Infrastructure for Integrated Structural Biology (FRISBI) ANR-10-INSB-05-01 and Instruct as part of the European Strategy Forum on Research Infrastructures (ESFRI).

## Methods (online methods section)

### Cell culture and immunofluorescence

For the tubulin and H2B samples, HeLa cells were cultured in a 4-compartment glass-bottom petri dish (CELLView, Greiner Bio-One), washed in PBS and fixed with 4% formaldehyde for 20 min in phosphate-buffered saline solution (PBS). After permeabilization with 0.1% Triton X-100 in PBS (PBS/Tx) twice for 10 min, the primary antibody (anti-β-tubulin monoclonal (1 Tub-2 A2, in house IGBMC) used as mouse ascites fluid diluted 500x in PBS/Tx; histone H2B monoclonal antibody (LG11-2) used as a 500x dilution of mouse ascites fluid in PBS/Tx) was incubated overnight at 4 °C. The sample was then washed with PBS/Tx three times over 30 minutes, and the secondary antibody (goat anti-mouse Alexa Fluor-647) in dilution 4 μg/ml in PBS/Tx was incubated for 1 to 2 hours at room temperature (RT). Subsequently, the cells were washed in PBS/Tx three times for 30 minutes, then briefly three times in PBS. The samples were mounted in a PBS buffer with addition of 10 mM of cysteamine (also known as β-mercaptoethylamine or MEA) and 25 mM of HEPES (pH 7.5).

For the CENP-A samples, U2OS cells were synchronized using Thymidine-Nocodazole synchronization. Briefly, the cells were plated and 24 hours later the medium was replaced with prewarmed complete DMEM supplemented with 2.5 mM Thymidine for 18 hours. The cells were washed with DMEM and 100 ng/ml Nocodazole was added for 12-14 hours. Floating cells were collected and spin down washed twice with prewarmed DMEM and resuspended in prewarmed DMEM. Cell were collected after 8 hours, washed twice in PBS and fixed in 2% paraformaldehyde for 15 min at RT, washed twice in PBS, permeabilized with PBS/Tx for 15 min, washed twice in PBS, washed five times in PBS+0.5% BSA (PBB), blocked with 10% serum in PBB for 1 hour, washed five times in PBB, incubated with primary antibodies (EMD Millipore 07-574) in 400x dilution for 1h, washed five times in PBB, incubated with secondary antibodies in 400x dilution for 1h, washed five times with PBB and three times with PBS. The sample was mounted in an imaging buffer^1^ that contained 20% of Vectashield (Vector Laboratories), 70% of 2,2-thiodiethanol (also known as thiodiglycol or TDE) and 10% of PBS 10x (the measured refractive index of this mounting medium is 1.49). The mounting was performed by incubating the sample in PBS solutions with gradually increased concentrations of TDE (10%, 25% and 50%), for 10 min each.

### Super-resolution imaging

The super-resolution imaging of the β-tubulin and the H2B samples was performed on a Leica SR GSD 3D system in Wetzlar, Germany. The system was equipped with the HC PL APO 160x/1.43 Oil CORR GSD objective and the Andor iXon Ultra 897 EMCCD camera with a field of view of 18x18 μm in the high-power mode. Continuous wave fiber laser (MPBC Inc., 642 nm 500 mW) and a diode laser (405 nm 30 mW) were utilized for excitation. The objective was linked to the sample with help of a suppressed motion (SuMo) sample stage, which reduces lateral and axial drift. The astigmatism was induced with the cylindrical lens and calibrated using gold beads and the automatic built-in procedures.

The super-resolution imaging of the CENP-A samples was performed on our in-house Leica SR GSD system. We used the HCX PL APO 100x/1.47 Oil CORR TIRF PIFOC objective with a 1.6x magnification lens that provides an equivalent pixel size of 100 nm on the camera. The camera, the lasers and the stage were the same as on the Leica SR GSD 3D system. The astigmatism was induced with a MicAO 3DSR adaptive optics system installed between the microscope stage and the camera (the value of the astigmatism was set to 0.2 μm root mean square). The deformation of the PSF was calibrated by defocusing the objective with a step of 50 nm and imaging Tetraspeck fluorescent beads with diameter of 200 nm.

The samples were first illuminated with the 100% power of the 642 nm laser to quickly send the fluorophores into the dark state. The acquisition was started manually after observing first single-fluorophore events (“blinking”). The time of exposition of a frame was 6.9 ms (H2B), 10 ms (β-tubulin) or 25 ms (CENP-A); the electron multiplying gain of the camera was 300; the laser power during the acquisition was 28% (H2B) or 100% (β-tubulin and CENP-A). After a few minutes, as the number of blinking evens dropped, the sample started to be illuminated additionally by a 405 nm laser with gradual increase of its intensity in order to keep a nearly constant rate of single-molecular returns into the ground state. The acquisition was stopped after almost complete bleaching of the fluorophore.

### Data processing

The localization and fitting of single-molecule events were performed in real time during acquisitions in Leica LAS AF software with the “direct fit” fitting method. The localization tables were then exported for further processing in the SharpViSu software workflow^2^ and with customized Matlab procedures. To reduce the number of localizations of the same fluorophore and improve localization precision the data were processed by averaging the coordinates of consecutive events within a radius of 50 nm around each localization. The drift was detected and corrected in three dimensions using cross-correlation-based approach. Briefly, the dataset was divided into several consecutive subsets, from each of them a histogram image with pixelation of 20 nm was build; next, the shift between these images was detected with subpixel precision and then interpolated linearly throughout intermediate frames. The shift value was then subtracted from the coordinates of every frame. The lateral drift was detected using the projection on the XY plane, and the axial drift was detected using the average shift, calculated from the XZ and YZ projections. The procedure was repeated iteratively several times assuring absence of detectable residual drift^2^. Regions of interest (ROI) for Voronoi analysis were selected manually allowing faster computations and more homogeneous distributions compared to the entire field of view.

The following analysis was performed in Matlab using customized code. Voronoi diagrams (vertices of polyhedrons and connectivity order) were retrieved with ‘voronoin’ function. Volumes of the cells were determined from the vertices with the function ‘convhulln’^3^, the local density in each data point was defined as the inverse value of the volume of the corresponding Voronoi polyhedron. To account for nonisotropic distribution of experimental data points in Z-direction we introduced a correction factor for the calculated local density. Indeed, the intensity of the PSF diminishes while going out of focus and so does the number of localization if one performs a 3D SMLM experiment using a PSF shape modifying method. Consequently, the local densities will be stronger in the center of the 3D volume compared to top and bottom regions in Z direction. To account for this distortion, we approximated the decreasing number of events in the axial direction as a Gaussian function with the fixed standard deviation σ = 300 nm, which corresponds to the axial resolution of the objective R_z_ = 1.5-λ-n/NA^2^, where λ is the wavelength of the detected light, n is the refractive index of the sample and NA is the numerical aperture of the objective (**Fig. S2**). The densities associated with the experimental localizations were then multiplied by a factor proportional to the inverse value of the Gaussian function at the Z-position of the fluorophore. To have a smooth appearance that can be used for visualization or segmentation the values of the corrected local densities were interpolated to a regular grid (voxels) using the ‘griddata’ function and the ‘natural’ interpolation method^4^ that is also based on Voronoi diagrams. The spacing of the grid corresponds to the desired voxel size (20 nm in our case).

For processing of experimental data acquired with astigmatism, the points for Monte-Carlo simulations were generated to have completely spatially random X, Y coordinates (the ‘rand’ function), and Z coordinates picked from a normal distribution with standard deviation σ = 300 nm (the ‘normrnd’ function). The points were distributed over the manually defined field of view (FOV). The distributions were generated for 50 different random sets of points, the boundaries of the confidence envelope were determined as <n> ± 1.96·σ for each bin of the histogram, where <n> is the average number of cells within the range of the bin and σ is the standard deviation of n, calculated from the 50 random datasets. The abscissas of the first and the second intersections between the curves of the experimental and the mean value of the randomized distributions were determined from the two points around the intersections in the linear approximation.

The simulated cluster data in **Figs. 2C-I** was generated as randomly distributed points inside spheres with the radius of 30 nm. The positions of clusters and of background points were distributed randomly in the FOV, with the following constraints: 1) distance from the centers of the clusters to the borders ≥ 75 nm; 2) distance between the centers of the clusters ≥ 150 nm. The borders of the clusters were smoothened such that the density of molecules at the distance r from the center of the cluster was modulated with the function 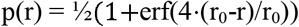, where r_0_ = 30 nm and erf(x) is the error function.

Even though the distribution of Voronoi cell areas of the 2D-projected dataset in Fig. 1C lies within the confidence envelope for the background model indicating no clustering, we segmented this dataset keeping only the regions with the local density stronger than the mean density calculated from the 2D Voronoi diagram^5^. The resulting clusters, as expected, have random size much smaller than the original radius of 30 nm (Fig. 1C, left and right panels).

We have also implemented the averaging of the local densities using the first-rank^5^ neighbors in 3D. This option helps to remove spurious clusters in the background but it also leads to reduction of the volume of the detected clusters because the density of the near-border regions within clusters is averaged with the low-dense background (**Fig. S3**). By contrast, simple removal of small clusters as done by default (Fig. 2H) does not affect the shape of big clusters. Although not done here, it is possible to separate closely located clusters through methods such as watershed transformation, which are compatible with our clustering method and can be performed after binarization of the density map. Similarly, if experimentally evidenced, the background noise model could be adjusted to include re-localization events arising from molecules blinking several times.

For the CENP-A data, after the segmentation, centromeres touching each other and small clusters in the background were removed from the analysis. To determine the size of the CENP-A clusters, 5 different ROIs with 200 well separate centromeres in total were analyzed. The equivalent radius (diameter) of clusters was calculated as the radius (diameter) of the sphere with the same volume as the clusters. Quantified properties of clusters (number of events, equivalent radius or diameter) are represented as mean ± standard deviation of the corresponding values. The density map can be readily exported to TIFF or MRC formats for further processing or visualization.

The standalone software for visualization of 3D Voronoi tessellations was written in Python using the Mayavi data visualization library^6^ along with NumPy^7^ and SciPy^8^. SMLM datasets are loaded as a 3D CSV point list file. The user can then navigate through the 3D Voronoi cells determined by the input points. The cells are colored according to their size, from blue (big cells) to red (small cells). The user can set a cell size threshold to see only smaller cells where events are aggregated, use a clipping box to see inside the cells and toggle the drawing of the event points. The size of the cells is computed in parallel using QHull's "ConvexHull" function^3^ from SciPy. Cells delimited by vertices outside of the event range are merely ignored. The Mayavi interface allows the user to change a lot of drawing parameters on-the-fly. Colors, opacity, orientations, lightning, clipping shape etc. are all tunable through the built-in parameter adjustment wizard.

As an example, it takes 6 hours to segment a large experimental dataset with a volume of ~200 μm^3^ that contains 10^6^ localizations using an i7 quad-core single processor with 32 Gb of memory. This is much faster in a single run than what could be done for example with the current 3D implementation of DBSCAN^9^ in VividSTORM^10^ where one single run takes 8 hours for 67 000 points (which is a rather small data set). In contrast, our Voronoi-based segmentation uses an automatically determined threshold value and takes in total 6 hours for 1 000 000 points (15 times as many) processed in one single run, i.e. it is more robust, non-dependent on user-set parameters and runs faster (in total range of 30-100 times faster depending on the number of trials needed and the complexity of the data). 3D DBSCAN was implemented in Matlab to test DBSCAN at different background noise densities (Fig. S1B-D). The volume of a cluster was calculated as the volume of the convex hull of the points comprising the corresponding cluster.

### Code availability

Matlab implementations of the 3D Voronoi segmentation method and the Python-based 3D Voronoi visualization software tools are available under: http://cbi-dev.igbmc.fr/cbi/voronoi3D/repository/archive.zip http://cbi-dev.igbmc.fr/cbi/voronoi3D

